# Enteric glial cells of the two plexi of the enteric nervous system exhibit phenotypic and functional inter- and intra-heterogeneity

**DOI:** 10.1101/2022.06.28.497986

**Authors:** Touvron Melissa, Wieland A. Bradley, Chloe L. Mariant, Hattenhauer R. Alex, Laurianne Van Landeghem

## Abstract

Enteric glial cells (EGC) are a prominent cell type of all layers of the gut wall, virtually controlling all gastrointestinal functions. While the development of transgenic mice has led to major advances in understanding EGC biology, *in vitro* models are still fairly limited and do not allow for the robust and reproducible establishment of primary cultures discriminating EGC from the inner *versus* outer layers of the gut wall. Here we report a novel method to separately grow EGC from the inner and outer layers of the intestinal wall from the same mouse with a high degree of purity and cell heterogeneity. Our results indicate that EGC from the inner layers of the gut wall exhibit higher calcium response to ATP when compared to EGC from the outer layers. We also show that inner EGC cultures express lower levels of the transcription factor Sox 10 as compared to outer EGC cultures, which mirrors *in situ* differential expression of Sox10 in submucosal (inner) *versus* myenteric (outer) plexus assessed using wholemounts. Confocal microscopy analyses of wholemounts further demonstrate that a majority of calretinin-expressing ganglionic cells of the submucosal plexus express the EGC marker S-100β, while this population is marginally represented in ganglia of the myenteric plexus. Altogether this study describes a novel method of EGC primary cultures permitting for the first time to compare inner *versus* outer EGC and provides *in vitro* and *ex vivo* evidence that inner EGC and outer EGC are phenotypically and functionally distinct.

## Introduction

The enteric nervous system (ENS) is defined as the third branch of the autonomic nervous system and corresponds to the gastrointestinal (GI) tract intrinsic nervous system. It is composed of enteric neurons and enteric glial cells (EGC), which derive from sacral and vagal enteric neural crest cells. While EGC are found in every layer of the gut wall, from the serosa to the mucosa, the bodies of enteric neurons are confined to ganglia, which interconnect to form a plexus. The mouse gut wall comprises two plexi, a first one located in the submucosa, called submucosal plexus, and a second one located between the circular and the longitudinal muscle layers, called myenteric plexus. While these two plexi functionally interact to regulate gut functions, it is widely accepted in the field that the myenteric plexus is predominantly involved in gut motility, while the submucosal plexus is the main regulator of mucosal functions, including barrier function and electrolyte secretion. In addition to these distinct functional roles, it is also known that the two ENS plexi harbor different types of neurons with specific functions (sensory neurons, interneurons, muscle motor neurons, secretomotor neurons, vasodilator neurons) associated with distinctive morphology, neurochemical coding, electrophysiological properties and target cell(s). In contrast, much less is known about potential phenotypical and functional differences between EGC of the two plexi.

Until fairly recently, EGC were merely considered as structural and trophic supportive cells to neurons and approached as a homogeneous cell population. Compelling evidence accumulating over the past two decades has now well-established that EGC interacts with all cellular components of the gut wall to directly and/or indirectly control each and every GI function. In addition, an elegant study from Boesmans and colleagues in 2015 has provided evidence that in wholemounts of myenteric plexus, EGC with discrete morphology and molecular identity exhibited distinct calcium response to ATP^1^, highlighting that myenteric EGC are a phenotypical and functional heterogeneous population. Further reinforcing this concept of functional heterogeneity, a recent paper from Baghdadi and colleagues identified that loss of GFAP-expressing EGC, but not PLP-1-expressing EGC, affects intestinal stem cell maintenance and crypt regeneration following injury, although the two EGC population can compensate for each other’s loss to maintain or restore intestinal homeostasis^2^. Collectively the literature strongly indicates that EGC are key cellular effectors and regulators of GI physiology and that the EGC network harbors distinct EGC subpopulations with specific phenotype and function^3^. However, the understanding of this cellular heterogeneity is far from being complete and the development of novel platforms is necessary to fill these gaps of knowledge.

In this work, we aimed to develop a simple, robust and reproducible method to grow separately EGC present in the most outer regions of the intestinal wall, *i*.*e*., EGC from the mucosa and submucosa, and EGC present in the most inner regions of the wall, *i*.*e*., EGC from the myenteric plexus, muscles and serosa. Although we acknowledge that all EGC from all layers most likely functionally cooperate together, we reasoned that such a tool would provide the opportunity to compare phenotypical and functional characteristics of primary cultures of EGC, either enriched in cells primarily involved in the regulation of mucosal/submucosal functions-that we termed inner EGC or inEGC - or enriched in cells primarily involved in the regulation of muscularis functions, *i*.*e*., gut motility -that we termed outer EGC or outEGC. Our data demonstrate that our culture conditions allow for the establishment of EGC primary cultures presenting a degree of purity and heterogeneity. Using this novel model, we uncover dramatic differences in calcium response to purinergic signaling and Sox10 expression between inEGC and outEGC. In addition, we use confocal analyses of wholemounts specimens to validate some of our findings *in situ* and we highlight additional fundamental phenotypical differences between inEGC and outEGC present in ganglia of respective plexus.

## Methods

### Animals

All animal studies were approved by the Institutional Animal Care and Use Committee of the North Carolina State University. CD1 mice were housed under standard conditions. Mice were anesthetized with isoflurane and sacrificed by cervical dislocation.

### Tissue collection and dissociation

The entire small intestine was collected, flushed with ice-cold 1X phosphate buffered saline (PBS) and divided into 1cm sections that were subsequently stored in cold 1X hanks balanced salt solution (HBSS). Individual sections were inserted onto a sterile 1,000µL tip. Serosa, longitudinal muscle and myenteric plexus along with circular muscle were gently stripped off from underlying layers with fine curved forceps and stored in cold 1X HBSS solution. Remnant tissues containing submucosa and mucosa were opened longitudinally and stored in cold 1X HBSS solution.

Tissues comprised of the submucosa and mucosa layers (inner layers) were first incubated in 10mL of cold 1X PBS containing 30mM EDTA pH8 and 1.5mM DTT for 15 minutes on ice and next transferred into 8mL of in 1X PBS with 30mM EDTA pH8 for 8 minutes at 37°C. Tissues were manually shaken to detach epithelial cells and remnant tissues were quickly washed by dipping them into 1X HBSS solution prior to incubation into 5mL of “dissociation” medium composed of DMEM/F12 medium (Dulbecco’s modified Eagle’s medium/F12, Gibco) supplemented with 1.19mg/mL NaHCO3 (Sigma), 2mM L-glutamine (Corning), 50U/mL Penicillin (GeminiBio), 50ug/mL Streptomycin (GeminiBio), 1.1ug/mL Amphotericin B (Sigma), 20ug/mL gentamycin (Sigma), 0.5mg/mL protease (Sigma), 0.25mg/mL collagenase (Sigma) and 4mg/mL Bovine Serum Albumin fraction V (Sigma). Tissues containing the serosa, the muscles and the myenteric plexus (outer layers) were also transferred into 5mL of “dissociation” medium. Enzymatic digestion of inner and outer layers was carried out for 7 and 15 minutes, respectively, at 37°C under constant gentle agitation. Tissues were next vortexed 3 times for 10 seconds and digestion was stopped by adding 10mL of DMEM/F12 medium (Gibco) supplemented with 1.19mg/mL NaHCO3 (Sigma), 2mM L-glutamine (Corning), 50U/mL Penicillin (GeminiBio), 50ug/mL Streptomycin (GeminiBio), 1.1ug/mL Amphotericin B (Sigma), 20ug/mL gentamycin (Sigma) and 10% heat-inactivated fetal bovine serum (FBS; GeminiBio). Tissues were next mechanically dissociated by pipetting up and down 15 times and vortexing 3 times for 10 seconds. After centrifugation for 5 minutes at 1,250rpm at room temperature (RT), pellets were reconstituted with 10mL of DMEM/F12 medium (Gibco) supplemented with 1.19mg/mL NaHCO3 (Sigma), 2mM L-glutamine (Corning), 50U/mL Penicillin (GeminiBio), 50ug/mL Streptomycin (GeminiBio), 1.1ug/mL Amphotericin B (Sigma), 20ug/mL gentamycin (Sigma) and 10% heat-inactivated fetal bovine serum (FBS; GeminiBio) and further dissociated by pipetting up and down 15 times and vortexing 3 times for 10 seconds. Cell preparations were sequentially filtered through 100µm, 70µm and 40µm cell strainers.

### EGC primary culture establishment and maintenance

Wells were coated with 0.5mg/ml poly-L-lysine (Sigma) overnight at RT then washed 2 times with double-distilled H2O. Cells from inner layers (inEGC) were plated at a density of 300,000 cells/well, while outer layers (outEGC) were plated at 100,000 cells/well in 24 well-plates. Cells were plated in DMEM/F12 (Gibco) supplemented with 10% heat-inactivated FBS (GeminiBio), 2mM L-glutamine (Corning), 50U/mL Penicillin (GeminiBio), 50ug/mL Streptomycin (GeminiBio), 1X G5 supplement (Gibco), 1X N2G supplement (GeminiBio) and 1X Gem21 supplement without vitamin A (GeminiBio) and switched to DMEM/F12 (Gibco) supplemented with 0.5% heat-inactivated FBS (GeminiBio), 2mM L-glutamine (Corning), 50U/mL Penicillin (GeminiBio), 50ug/mL Streptomycin (GeminiBio), 1X G5 supplement (Gibco), 1X N2G supplement (GeminiBio) and 1X Gem21 supplement without vitamin A (GeminiBio) the next day. Medium was changed every 2-3 days. EGC primary cultures were passaged when cultures reached 60-70% confluence or when 3D clusters started to form using Trypsin-EDTA 0.25% (GeminiBio) diluted 1:5 in 1X PBS. Investigations presented here were carried out using inEGC and ouEGC primary cultures at passage 0 (P0) and passage 1 (P1).

Purity and cellular heterogeneity were assessed by immunofluorescence in P0 and P1 EGC primary cultures grown in 24 well-plates and fixed for 15 minutes in 4% paraformaldehyde in 1X PBS at RT. Calcium assays were conducted on P0 EGC primary cultures grown in 96 well plate.

### Wholemount dissection

Small intestine was collected, flushed with ice-cold 1X PBS and cut open along the mesentery. A ∼ 3cm long section was gently pinned and stretched on Sylgard-coated glass dishes with the mucosal side facing up. Pinned specimens were next fixed in 4% paraformaldehyde in 1X PBS overnight at 4°C.

With fine micro-dissection forceps, the mucosa layer was removed. The submucosal layer containing the submucosal plexus was carefully isolated using microscissors and fine curved forceps. Circular muscle fibers were gently stripped off using fine straight forceps to isolate the myenteric plexus resting on the longitudinal muscle layer covered by a thin serosa, commonly called LMMP. 25mm square specimens of submucosa and LMMP were stored in 1X PBS with 0.1% NaN3 at 4°C until being processed for staining.

### Immunofluorescence staining for 2D imaging and confocal microscopy

EGC primary cultures and wholemount tissues were incubated in 1X PBS containing 5% Horse Serum and 0.1% Triton-X-100 for 30 minutes at RT prior to incubation with primary antibodies diluted in 1X PBS containing 5% Horse Serum and 0.1% Triton-X-100 for 3 hours at RT or overnight at 4°C, respectively. The list of primary antibodies used in this study are shown in Supplemental Table 1. After extensive washes in 1X PBS, cells and wholemounts were next incubated with the secondary antibodies diluted in 1X PBS containing 5% Horse Serum and 0.1% Triton-X-100 for 45 minutes in the dark at RT. The list of secondary antibodies used in this study are shown in Supplemental Table 1. Different sets of primary/secondary antibodies were incubated sequentially to avoid cross-reactions. Following extensive washes in 1X PBS, nuclei of primary cultures were counterstained with 5ug/ml DAPI for 5 minutes at RT, while wholemount tissues were mounted under dissecting microscope between slide and glass coverslip using a mounting gel with DAPI (Electron Microscopy Sciences).

EGC primary cultures were imaged using an Olympus IX83 inverted microscope equipped with an ORCA-Flash4.0 V2 camera from Hamamatsu using a 20X objective (LWD U PLAN FL PHI, NA 0.45, .4-7.6MM) and images were analyzed using imageJ. Purity and heterogeneity were assessed until 1,000 cells were analyzed from multiple randomly selected representative images or on entire well if well contained less than 1,000 cells.

Wholemount tissues were imaged using a Nikon A1R confocal microscope equipped with a Hamamatsu camera using a 20X objective (Plan Apo VC DIC, NA 0.75) and images were analyzed using imageJ.

### Immunofluorescence staining for light sheet microscopy using the iDISCO+ procedure

Full thickness small intestines were pinned onto Sylgard-coated glass dishes and fixed in 4% paraformaldehyde overnight at 4°C. After extensive 1X PBS washes, samples were dehydrated, bleached, immunostained and clarified according to the iDISCO protocol described by Renier and colleagues. Briefly, samples were dehydrated for 1 hour at RT in ascending concentrations of methanol (20%, 40%, 60%, 80%, 100%) and incubated overnight at RT in 66% dichloromethane-33% methanol. After methanol washes, tissues were bleached in a 5% hydrogen peroxide solution-100% methanol overnight at 4°C and re-hydrated on the following day in descending concentrations of methanol (80%, 60%, 40%, 20%) followed by 1X PBS washes. Samples were permeabilized in 1X PBS containing 23 g/L glycine (Sigma-Aldrich), 0.2% Triton X-100 (Sigma-Aldrich) and 20% DMSO (Sigma-Aldrich) for 2 days at 37°C and blocked in 1X PBS containing 0.2% Triton X-100, 10% DMSO and 6% donkey serum (Jackson Immunoresearch) for 2 days at 37°C. Immunostaining for S-100β (DAKO, Z0311, 1:500) was performed in 1X PBS containing 0.2% Tween 20 (Sigma-Aldrich), 10 µg/ml Heparin (Sigma-Aldrich), 5% DMSO and 3% donkey serum at 37°C for 4 days, followed by 5 washes of 1 hour in 1X PBS with 0.2% Tween 20 and 10 µg/ml Heparin at RT. Tissues were then incubated for 4 days at 37°C with a secondary antibody (donkey anti-rabbit AF488, Invitrogen, 1:1,000) in 1X PBS with 0.2% Tween 20, 10 µg/ml Heparin and 3% donkey serum. After washes in 1X PBS with 0.2% Tween 20 and 10 µg/ml Heparin at RT, tissues were mounted in 1% agarose (Invitrogen) diluted in 1X TAE (Tris-Acetate-EDTA buffer) and left in 0.2% Tween 20 and 10 µg/ml Heparin at RT overnight. The next day, samples were dehydrated in ascending concentrations of methanol (20%, 40%, 60%, 80%, 100%) for 1 hour at RT and left overnight in 100% methanol. Samples were then cleared in 66% dichloromethane-33% methanol for 3 h at RT and then washed in dichloromethane. Samples were subsequently transferred and stored in 100% dibenzyl ether solution (DBE, Sigma-Aldrich). Samples were imaged using a LaVision BioTec UltraMicroscope II light sheet system equipped with zoom body optics, an Olympus MV PLAPO 2XC 2X/0.5 objective, and a LaVision BioTec corrected DBE dipping cap with a 5.7 mm working distance. The zoom setting was always 1.6X, for a combined magnification of 3.2X, corresponding to a 1.91 µm pixel size. Samples were imaged sequentially from both sides, with sheet NA set to 0.026 (estimated Z thickness at waist = 28 µm), and sheet waist centered on the midpoint of each side of the sample. LaVision’s InspectorPro software automatically combined the two images into one by merging the left side of the image illuminated with the left light sheet, and the right side of the right image illuminated with the right light sheet, with a small interpolation region in the middle. Images were acquired by sequentially taking Z stacks with 5 µm Z spacing. Images were generated with Imaris 9.2 (Bitplane) software by cropping in 3D to trim away rough borders of the sample and adjusting display settings.

### Calcium activity assay

Calcium response to ATP was assessed using the Fluo-4 calcium imaging kit (Life technologies) following the manufacturer’s recommendations. Live-cell imaging was performed using an Olympus IX83 inverted microscope equipped with an ORCA-Flash4.0 V2 camera from Hamamatsu using a 20X objective (LWD U PLAN FL PHI, NA 0.45, .4-7.6MM). Basal fluorescence of each well was recorded for 60 seconds. Subsequently, changes in fluorescence were recorded for 60 seconds following addition of ATP (Sigma) at 0.5mM, 2mM or 10mM to the well. Movies were transformed into image sequences using Photoshop (Adobe) at 12 frames/second, and whole-field fluorescence intensity was determined for each frame/image using image J. Graphs f(time) = (fluorescence intensity) were created using Excel (Microsoft) and areas under the curve were calculated using ImageJ. Area under the curve calculated from the recording of basal fluorescence was subtracted for each well to account for potential background noise. Data were next normalized to the surface covered by EGC in each well.

### Statistics

Data were expressed as means ± SEM. Statistical analyses were performed using SigmaPlot and GraphPad Prism software. Non-parametric tests were used to compare different groups as indicated in the figure legends if Shapiro-Wilk Normality Test or Equal Variance Test failed. Differences were considered as significant for a p-value of less than 0.05.

## Results

### Novel methods of isolation and culture allow for the establishment of primary cultures of EGC originating from the outer versus the inner wall layers of the small intestine

Most published methods for the establishment of EGC primary cultures are either based on the isolation of one plexus only^4–7^ or do not discriminate the two plexi of the ENS^8^. However, the submucosal plexus and the myenteric plexus are morphologically distinct as illustrated in Figure 1 in which we show full thickness sections of adult mouse small intestines cleared, stained for the EGC marker S-100β and imaged using light sheet microscopy following the iDISCO method^9^. Marked differences between the two ENS plexi are readily observable, with the submucosal plexus exhibiting smaller ganglia and thinner inter-ganglionic fibers associated with a reduced EGC density as compared to the myenteric plexus (Figure 1 and Video 1). We thus sought to develop a method to discriminately isolate and grow EGC from the outer layers of the gut wall (outEGC) and EGC from the inner layers of the gut wall (inEGC) in the adult mouse. To do this, outEGC were isolated by peeling off the muscle layers enclosing the myenteric plexus, while inEGC were obtained following an EDTA/DTT treatment of the remnant tissue combined with mechanical dissociation to detach the epithelium from the lamina propria (Supplemental Figure 1a-c). Cellular preparations were next enzymatically digested and mechanically dissociated to obtain single cell preparations plated on poly-L-lysine and grown in reduced-serum (0.5%) medium enriched with N2, B27 and G5 supplements (Supplemental Figure 1d-e). Resulting cultures were analyzed at DIV4 (day *in vitro*). Primary cultures of outEGC and inEGC were composed of cells with similar average sizes (Figure 2a-b) and showed a similar distribution of cells with different sizes (Supplemental Figure 2a). Of note, while both outEGC and inEGC cultures formed clusters of cells at later time points of cultures (>DIV7-8), outEGC consistently grew clusters of cells at early time points (DIV3/4) (Figure 2a). To evaluate the purity of outEGC and inEGC primary cultures, we co-stained the cultures using antibodies against S-100β, PGP9.5 and αSMA to identify EGC, enteric neurons and myofibroblasts/smooth muscle cells, respectively (Figure 2c). Analyses showed that outEGC and inEGC cultures were composed of >80% of cells S-100β^+^/PGP9.5^-^/αSMA^-^ (Figure 2d). Additional staining showed that most cells present inEGC and outEGC primary cultures were strongly immunoreactive for the p75 neurotrophin receptor (Supplemental Figure 2b). We observed that the remaining ∼20% of cells were either immunoreactive for PGP9.5, αSMA or negative for all 3 markers (S-100β^-^/PGP9.5^-^/αSMA^-^). Of note, the majority of (if not, all) cells positive for PGP9.5 or αSMA were also positive for S-100β, but no cell was found to express both PGP9.5 and αSMA. Additional staining suggested that inEGC and outEGC primary cultures contained rare cells positive for F4/80, a marker of macrophages (Supplemental Figure 2c). Passaging of primary cultures did not significantly affect the purity of inEGC and outEGC primary cultures (Figure 2c-d and Supplemental Figure 2c). Interestingly P1 outEGC were significantly larger than P0 outEGC (941.7µm^2^ ±95.7 *vs*. 653.1 µm^2^ ±48.3, Mann-Whitney, n=4, p<0.05), but we found no statistical differences between P1 inEGC and P0 inEGC (863.6µm^2^ ±39.1 *vs*. 691.3 µm^2^ ±80.7, Mann-Whitney, n=4). Together these data validate our methods to establish primary cultures of EGC from the outer *versus* inner layers of the small intestine wall, which allow, for the first time, for phenotypical and functional comparisons between the inEGC *versus* outEGC.

**Figure 1.**
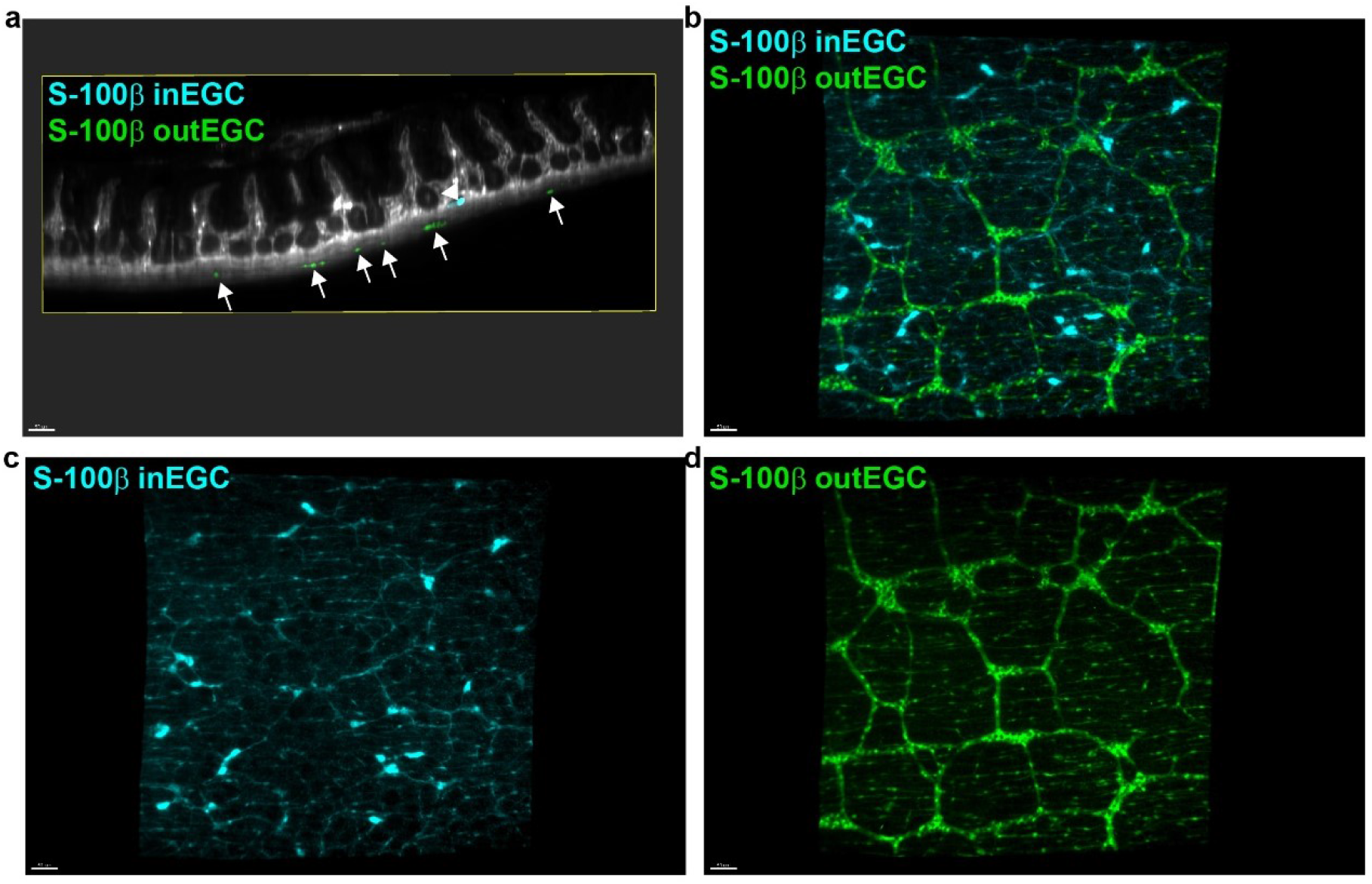
The two plexi of the ENS are morphologically distinct. 3D imaging of an adult mouse ileum using light sheet microscopy after clearing and staining for the EGC marker S-100β using the iDISCO (immunolabeling-enabled three-dimensional imaging of solvent-cleared organs) method. (a) x-z orthogonal projection allowing for the visualization of the 2 ENS plexi using the S-100β signal (teal and green). The signals originating from the submucosal plexus (**inEGC; teal; arrowhead**) and from the myenteric plexus (**outEGC; green; arrows**) are shown in teal and green, respectively. (b-d) x-y maximal projection showing that the two ENS plexi (**inEGC; teal** *vs*. **outEGC; green**) exhibit profound architectural differences. Scale bars = 50 µm.

**Figure 2.**
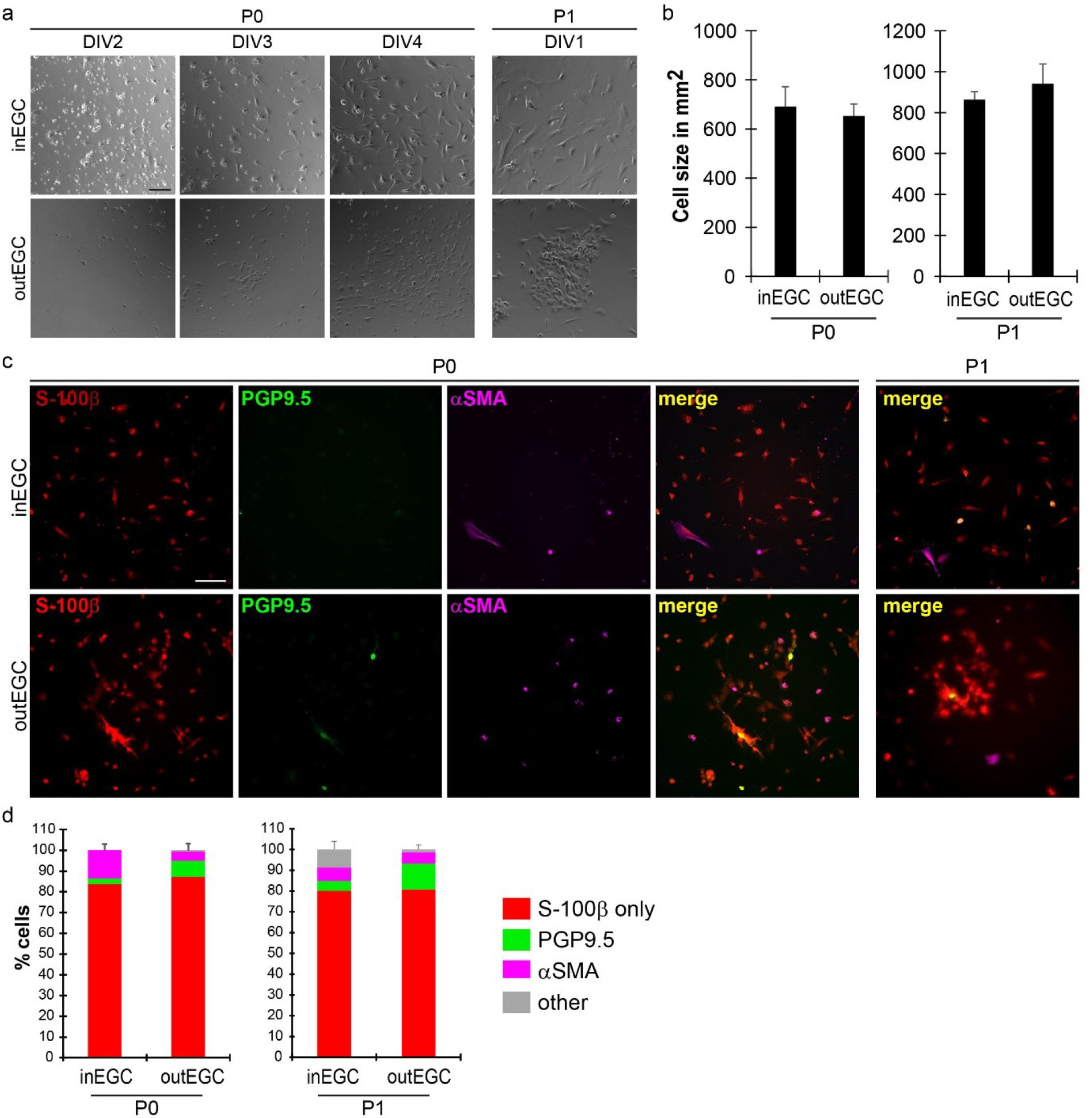
Primary cultures of inEGC and outEGC show similar cell sizes and high purity. (a) Representative brightfield images of P0 (passage 0) and P1 (passage 1) inEGC and outEGC primary cultures at DIV1-4. (b) Bar graphs showing that P0 and P1 inEGC and outEGC exhibit similar average cell sizes. 61±3 cells have been sized per condition. n=4 independent experiments. (c) Representative immunofluorescence pictures of P0 and P1 inEGC and outEGC primary cultures stained for S-100β (EGC marker), PGP9.5 (pan-neuronal marker) and αSMA (myofibroblast/smooth muscle cell marker). (d) Quantification of the percentage of cells immunoreactive for S-100β only (S-100β^+^/PGP9.5^-^/αSMA^-^), PGP9.5, αSMA or negative for all 3 markers (other; S-100β^-^/PGP9.5^-^/αSMA^-^). Of note, a vast majority of (if not all) cells positive for PGP9.5 or αSMA were also positive for S-100β, but no co-localization was observed between PGP9.5 and αSMA. 638±147 cells have been sized per condition. n=4 independent experiments. Scale bars = 100µm.

### Primary cultures of EGC from the outer versus the inner layers of the gut wall show distinct compositions of different EGC sub-populations

We next sought to determine whether inEGC and outEGC primary cultures were composed of different subtypes of EGC. To do this, we co-stained inEGC and outEGC cultures with antibodies against GFAP, S-100β and Sox10, all markers of EGC. Confirming the data shown in Figure 2, a vast majority of cells were immunoreactive for S-100β in both inEGC and outEGC primary cultures (Figure 3a). In addition, we observed that almost all inEGC and outEGC were positive for GFAP (Figure 3a). However, we noticed a striking difference in Sox10 immunoreactivity between inEGC and outEGC, with inEGC showing only sparse cells with weak immunoreactivity for Sox10 as compared to outEGC which exhibited a high proportion of cells strongly expressing Sox10 (Figure 3a). Thorough analyses indicated that cells in inEGC and outEGC cultures were expressing GFAP, S-100β and Sox10 at discrete levels of expression as illustrated in Figure 3b. For each marker, two levels of expression were easily discriminated, and we thus qualified cells as “high” when they exhibited extremely bright fluorescence or as “low” when they exhibited moderate fluorescence, yet significantly stronger than negative cells or background. To determine whether inEGC and outEGC cultures showed differences in the proportions of cells with high or low levels of expression of GFAP, S-100β and Sox10, we evaluated the number of cells with high or low levels of GFAP, S-100β and Sox10 independently in entire wells of inEGC and outEGC primary cultures (Figure 3b). Our analyses showed that P0 inEGC showed significantly less cells positive for Sox10 as compared to outEGC and that this was due to a lower proportion of cells expressing high levels of Sox10 (Figure 3b). A majority of cells were found to express relatively low levels of S-100β in both P0 inEGC and outEGC (Figure 3b). While P0 outEGC showed approximately equal proportions of cells with low and high levels of GFAP, P0 inEGC cultures were composed by a majority of cells with low level of GFAP (Figure 3b). Passaging of primary cultures did not significantly affect the relative proportions of S-100β high and low cells in both inEGC and outEGC (Figure 3b). Similar to P0 cultures, P1 outEGC had significantly a higher proportion of Sox10 positive cells due to a higher proportion of Sox10 high cells as compared to P1 inEGC. In fact, 90.9%±13.8 Sox10 positive cells in P1 outEGC cultures were Sox10 high cells. In addition, the differences in the proportion of GFAP high cells observed between P0 inEGC and P0 outEGC were abolished with passaging, with P1 inEGC and P1 outEGC both comprising approximately 60% of GFAP high cells (Figure 3b). We next quantified the proportions of cells co-expressing high or low levels of GFAP, S-100β and Sox10 to further determine whether inEGC and outEGC cultures had distinct compositions of EGC subtypes. We thus defined 13 EGC subtypes as shown in Figure 3c. Our analyses indicated that the main subtype in both P0 inEGC and P0 outEGC was GFAP low/S-100β low/Sox10 negative cells (subtype #12), although inEGC cultures had significantly more GFAP low/S-100β low/Sox10 negative cells (subtype #12) as compared to outEGC (69.8%±1.7 *vs*. 39.9%±8.4, Mann-Whitney, n=4, p<0.05) (Figure 3c). The major difference between P0 inEGC and P0 outEGC compositions was that P0 inEGC were devoid of Sox10 high cells (Figure 3c). In addition, amongst the 4 subtypes of P0 outEGC with high levels of Sox10, cells with a high level of GFAP and a high level of S-100β were significantly predominant (22.7%±6.5 (subtype #1) *vs*. 0.3%±0.1 (subtype #2), 2.4%±2.0 (subtype #3), 0.1%±0.1 (subtype #4), ANOVA, n=4, p<0.05). Passaging of primary cultures only induced subtle changes in the composition of inEGC and outEGC primary cultures (Figure 3c). Specifically the predominant subtype in P1 inEGC changed to GFAP high/S-100β low/Sox10 negative cells (subtype #11). In addition, amongst the 4 subtypes of P1 outEGC with high levels of Sox10, the predominant subtype changed to cells with a high level of GFAP and a low level of S-100β (30.0%±5.8 (subtype #3) *vs*. 8.2%±2.9 (subtype #1), 0.4%±0.2 (subtype #2), 2.9%±0.8 (subtype #4), ANOVA, n=4, p<0.05). Taken together our data indicate that inEGC and outEGC primary cultures are heterogeneous cultures since they comprise different subtypes of EGC based on GFAP, S-100β and Sox10 expression. Our data further show that inEGC and outEGC show distinct compositions of the different EGC subtypes.

**Figure 3.**
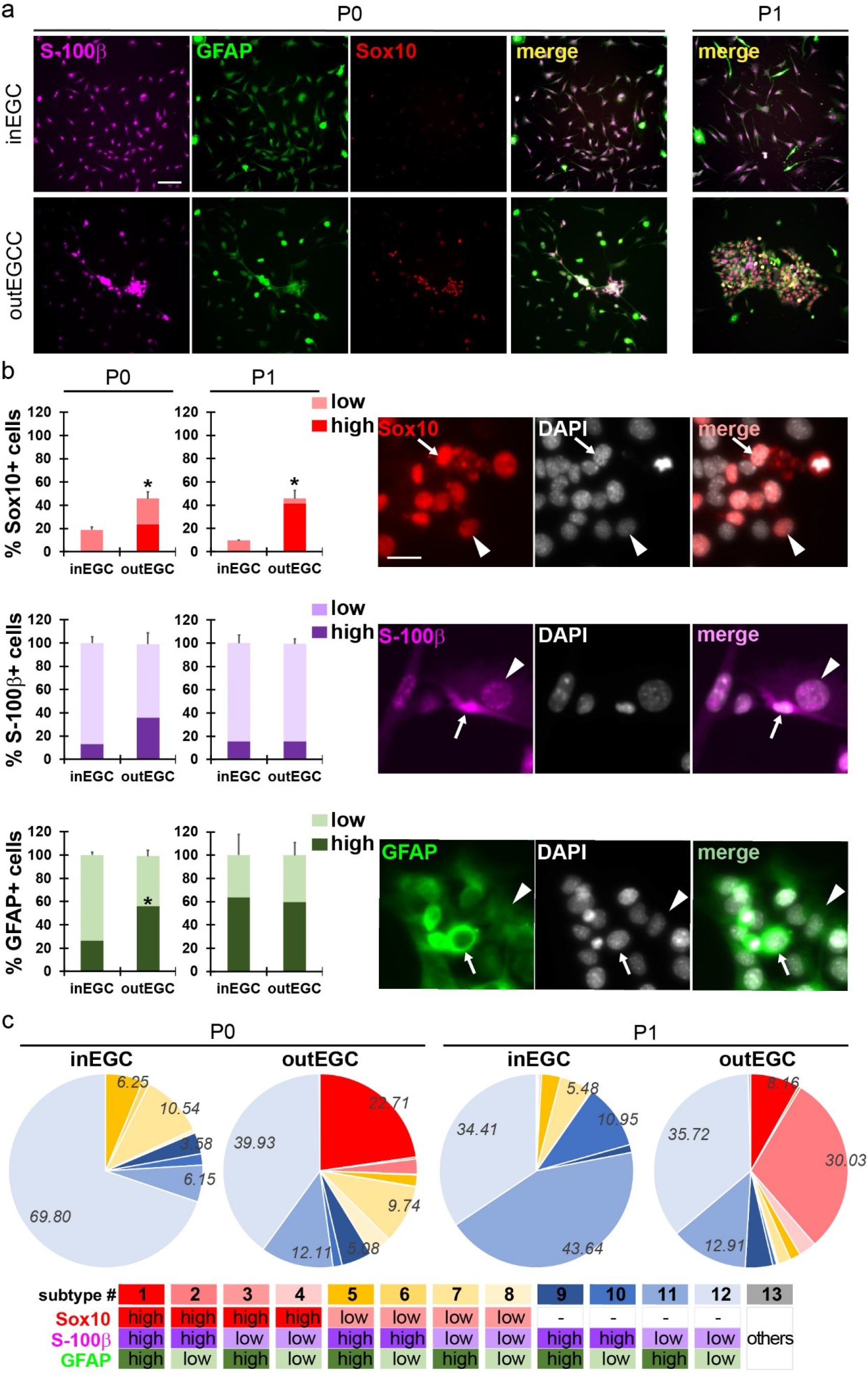
Primary cultures of inEGC and outEGC show distinct distributions of EGC sub-populations. (a) Representative images of P0 (passage 0) and P1 (passage 1) inEGC and outEGC primary cultures stained for S-100β (Magenta), GFAP (green) and Sox10 (red) demonstrating that a great majority of cells are positive for GFAP and S-100β in inEGC and outEGC cultures, and that while outEGC exhibit multiple cells (strongly) expressing Sox10, inEGC show a very small number of cells (weakly) expressing Sox10. Scale bar = 100µm(b) Bar graphs representing the proportion of cells in P0 and P1 inEGC and outEGC primary cultures showing high or low immunoreactivity for Sox10 (red), S-100β (Magenta) and GFAP (green). Data are expressed as the average of the percentages calculated per well (±SEM) (entire wells were analyzed). n=4 independent experiments. Mann-Whitney, *p<0.05. Representative images have arrow heads to show cells with low immunoreactivity, and arrows to show cells with high immunoreactivity. Scale bar = 20µm. (c) Pie charts representing the relative distribution of the different 13 subtypes of EGC in P0 and P1 inEGC and outEGC primary cultures based on the co-expression of low or high levels of Sox10, S-100β and GFAP. Thirteen subtypes are defined in table. Data are expressed as the average of the percentages calculated per well (±SEM) (entire wells were analyzed). n=4 independent experiments.

### Primary cultures of EGC from the inner layers of the gut wall show higher calcium response to ATP as compared to primary cultures of EGC from the outer layers of the gut wall

Biological activity of glial cells, including EGC, is largely regulated by intracellular calcium signaling^10^. Glial calcium responses are triggered by the activation of cell surface receptors leading to changes in concentration of intracellular calcium. Amongst the many extracellular cues evoking calcium mobilization in EGC, ATP has been shown to be a broad and potent activator of myenteric EGC (comprised in outEGC)^11^. While less is known about submucosal EGC (part of inEGC) response to ATP, recent studies suggest that purinergic signaling triggers intracellular transients in submucosal EGC^12^. To investigate whether inEGC and outEGC exhibited functional differences, we chose to measure changes in intracellular calcium concentration in P0 inEGC and outEGC primary cultures in response to 0.5mM, 2mM and 10mM ATP (Figure 4a-b). Our data showed that 2mM ATP induced the strongest calcium responses in both inEGC and outEGC primary cultures (Figure 4b). Importantly we found that inEGC primary cultures showed increased calcium responses when compared to outEGC (Figure 4a-b). While this was observed at all ATP concentrations tested, this was only statistically significant at 2mM ATP (6.9±2.1 *vs*. 1.4±0.6 arbitrary units, Mann-Whitney, n=6, p<0.05). In light of these data, we sought to determine whether this difference in ATP-evoked calcium responses between inEGC and outEGC was specific to selective subtypes of EGC. To do this, we analyzed calcium responses to 2mM ATP in single cells from each of the subtypes defined in Figure 3c -based on GFAP, S-100β and Sox10 expression- (Figure 4c). Our analyses showed that overall all subtypes of inEGC exhibited increased calcium signaling in response to ATP as compared to outEGC, and this was in particular significant for subtypes #12, #7 and #8 (Figure 4c). In addition, to test if differential levels of Sox10 expression were responsible for distinct calcium responses to ATP, we compared calcium responses between inEGC subtypes (ANOVA, Tukey’s multiple comparison test) and between outEGC subtypes (ANOVA, Tukey’s multiple comparison test). Analyses showed that there were no significant differences in calcium responses to ATP across all inEGC subtypes or all outEGC subtypes. Altogether these data demonstrate that inEGC have increased calcium signaling in response to purinergic stimulation as compared to outEGC and that this is independent from their levels of expression of Sox10, S-100b and GFAP.

**Figure 4.**
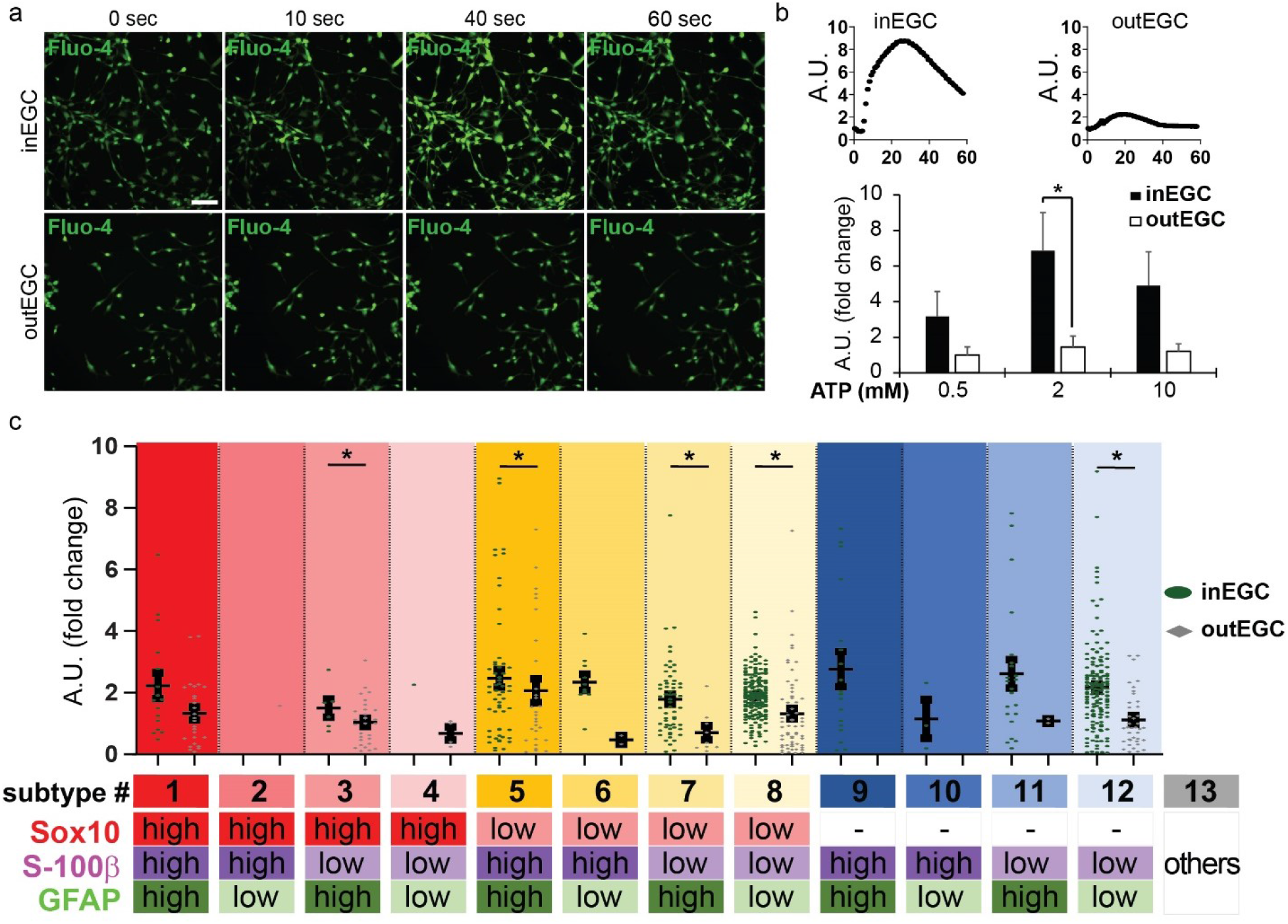
Primary cultures of inEGC and outEGC exhibit differential degrees of calcium response following ATP stimulation. (a) Representative sequences of images of P0 (passage 0) inEGC and outEGC primary cultures loaded with Fluo-4 (green) and stimulated with 2mM ATP. Scale bar = 100 µm. (b) *upper panel* Representative calcium responses recorded in inEGC and outEGC primary cultures stimulated with 2mM ATP. *lower panel* Graph represents intensity of calcium responses to 0.5mM, 2mM and 10mM ATP recorded in inEGC and outEGC primary cultures. Data are expressed as the averages of calcium response fold changes relative to the calcium response recorded in outEGC (established from the same animal) treated with 0.5mM. (A.U.C: area under the curve) (±SEM). n= 6 independent experiments (3 regions of interest were recorded for each dose in each experiment). Mann-Whitney, *p<0.05. (c) Graph represents intensity of calcium responses to 2mM ATP recorded in single cells from inEGC and outEGC primary cultures classified according to the 13 subtypes defined by the co-expression of low or high levels of Sox10, S-100β and GFAP. Data are expressed as calcium response fold changes relative to the average calcium response recorded in (all 13 subtypes of) outEGC (established from the same animal) (±SEM). n=0-180 cells were evaluated per subtype from 4 independent experiments (individual cells are shown on graph). Mann-Whitney, *p<0.05.

### Differences in Sox10 expression between inEGC and outEGC are observable in wholemount preparations of submucosal and myenteric plexi

To achieve the dual goal of demonstrating (1) that phenotypical differences exist *in vivo* between EGC from inner *versus* outer regions of the gut wall and (2) that our novel culture system faithfully recapitulates these differences, we sought to investigate whether Sox10 expression differed between EGC of the submucosal plexus -which are comprised in inEGC primary cultures- and EGC of the myenteric plexus -which are comprised in outEGC primary cultures-using wholemount preparations. Our data showed that the submucosal plexus presented a marked reduced number of Sox10-expressing cells per ganglion area as compared to the myenteric plexus (Figure 5a-b). Consistent with this result, we also found a decrease in the proportion of EGC expressing Sox10 in the submucosal plexus -assessed by normalizing the number of Sox10 positive cells to S-100β intensity per ganglion area (Figure 5a, c). These results confirm our hypothesis that inEGC and outEGC present major phenotypical differences that are captured by our *in vitro* culture system.

**Figure 5.**
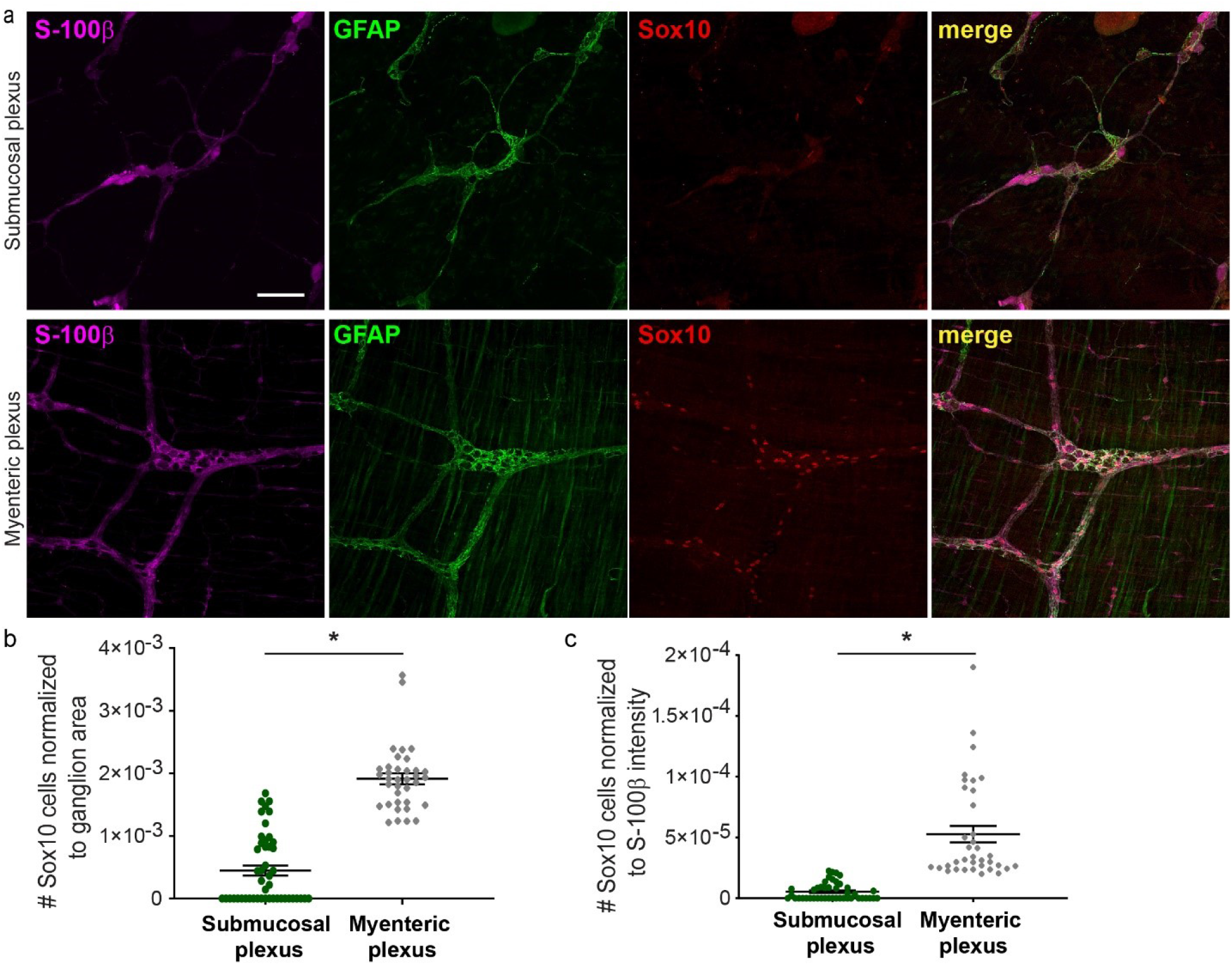
Myenteric plexus exhibits increased proportion of EGC expressing Sox10 as compared to submucosal plexus in wholemount preparations. (a) Representative confocal images of wholemount preparations of ileal submucosal and myenteric plexi from an adult mouse stained for S-100β (magenta), GFAP (green) and Sox10 (red). Scale bar = 100µm. (b) Quantification of the number of Sox10-expressing cells normalized to the ganglion area in submucosal plexus *vs*. myenteric plexus. Graph shows individual ganglia (n=36-47 ganglia) from 3 independent experiments. Mann-Whitney, *p<0.05. (c) Quantification of the number of Sox10-expressing cells normalized to S-100β intensity in the region of interest (i.e., ganglion) in submucosal plexus *vs*. myenteric plexus. Graph shows individual ganglia (n=36-44 ganglia) from 3 independent experiments. Mann-Whitney, *p<0.05.

### The majority of calretinin-expressing submucosal neurons are immunoreactive for S-100β

While characterizing S-100β immunoreactivity in wholemount preparations of submucosal and myenteric plexi, we noticed that larger S-100β-positive ganglionic cells were readily observable in the submucosal plexus, but not in the myenteric plexus (Figure 5a). To investigate this additional phenotypic difference between the two ENS plexi, we attempted to define the cellular identity of these large S-100β-positive cells. Due to their morphology and position within the ganglia, we hypothesized that these cells may correspond to previously described enteric neurons. To test this hypothesis, we co-stained wholemount preparations of submucosal and myenteric plexi for the neuronal marker PGP9.5 and S-100β (Figure 6a) and quantified the proportion of PGP9.5-expressing cells immunoreactive for S-100β (Figure 6c). Our data demonstrate that a majority of PGP9.5 cells are immunoreactive for S-100β in submucosal ganglia (56.5%±1.9), while only rare PGP9.5 cells were found to be S-100β-positive in myenteric ganglia (11.6%±0.9) (Figure 6a, c). We recapitulated these results using a second pan-neuronal marker, HuC/D (Figure 6 b, d). The dual expression of neuronal and glial markers within the same ganglionic cell was restricted to S-100β, as we did not find any PGP9.5 or HuC/D-expressing ganglionic cells immunoreactive for the EGC marker GFAP in both the submucosal and myenteric plexus (Figure 6 e-f). Co-localization of S-100β and GFAP in submucosal and myenteric ganglia further confirms that the large S-100β-positive ganglionic cells are GFAP negative and are predominantly found in the submucosal plexus (Figure 6g).

**Figure 6.**
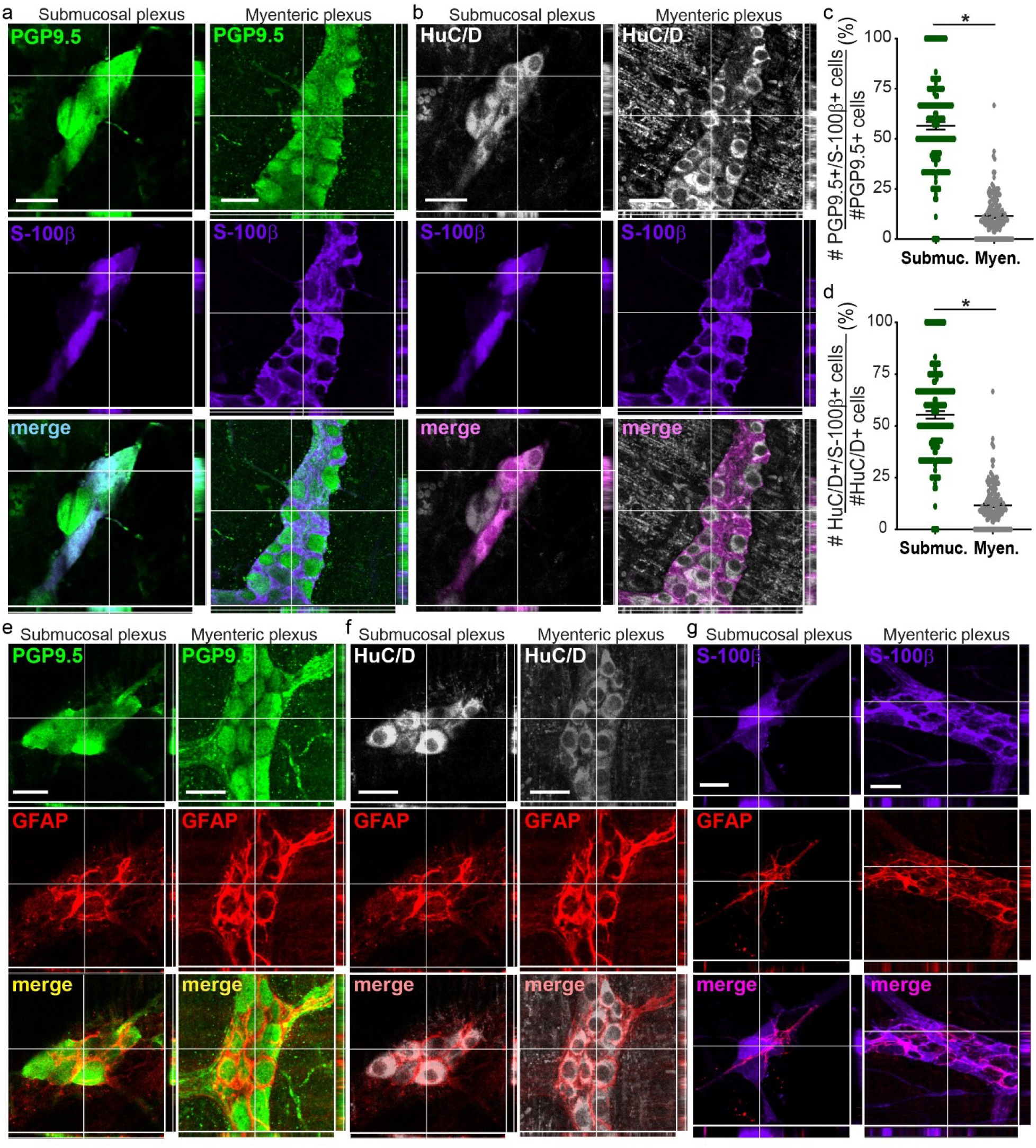
Ganglionic cells expressing both neuronal markers and S-100β are predominantly found in the submucosal plexus. (a-b) Representative confocal images of ganglia from wholemount preparations of ileal submucosal and myenteric plexi from an adult mouse stained for the neuronal markers PGP9.5 (a) (green) and HuC/D (b) (white) and the EGC marker S-100β (magenta). Scale bar = 30µm. (c-d) Quantification of the proportion of PGP9.5 (c) or HuC/D (d)-expressing cells immunoreactive for S-100β. Data are expressed as the percentages of the number of PGP9.5^+^/ S-100β^+^ cells normalized to the total number of PGP9.5^+^ cells per ganglion (c) and the percentages of the number of HuC/D^+^/ S-100β^+^ cells normalized to the total number of HuC/D^+^ cells per ganglion (d) (±SEM). Graphs show individual ganglia (n=130-153 ganglia) from 3-4 independent experiments. Mann-Whitney, *p<0.05. (e-f) Representative confocal images of ganglia from wholemount preparations of ileal submucosal and myenteric plexi from an adult mouse stained for the neuronal markers PGP9.5 (e) (green) and HuC/D (f) (white) and the EGC marker GFAP (red). Scale bar = 30µm. (g) Representative confocal images of ganglia from wholemount preparations of ileal submucosal and myenteric plexi from an adult mouse stained for S-100β (magenta) and GFAP (red). Scale bar = 30µm.

To further define the cellular identity of the PGP9.5^+^/S-100β^+^ ganglionic cells, we co-stained wholemount preparations of submucosal and myenteric plexi for S-100β and calretinin, which expression has been reported to be predominant in submucosal neurons^13^ and in some intrinsic sensory neurons, interneurons and excitatory motor neurons in the myenteric plexus^14,15^. We also co-stained wholemount preparations for S-100β and nNOS, which is known to be expressed in interneurons and inhibitory motor neurons in the myenteric plexus^14^, and which expression in the mouse ileal submucosal plexus is unclear^16– 18^. Our data show that over half (51.0%±6.2) of the calretinin-expressing submucosal neurons were immunoreactive for S-100β, while only a few (15.4%±1.5) calretinin-expressing myenteric neurons were positive for S-100β (Figure 7a-b). Consistent with this result, ganglia of the submucosal plexus exhibited a higher density in CAL^+^/S-100β^+^ cells as compared to the myenteric plexus (Figure 7c), despite ganglia of the myenteric plexus having the highest density of CAL^+^ neurons (Supplemental Figure 3a). We found that no submucosal ganglion showed co-localization nNOS and S-100β (Figure 7d-f), and this was due to an absence of nNOS immunoreactivity in ganglionic cells in the submucosal plexus, which is consistent with previous studies demonstrating <4% submucosal neurons were immunoreactive for nNOS^19,20^ (Supplemental Figure 3b). In addition, while nNOS-expressing neurons were readily observable in myenteric ganglia (Figure 7d and Supplemental Figure 3b), only a few a few (15.7%±2.9) nNOS-expressing myenteric neurons were positive for S-100β (Figure 7e). Altogether these data show that a majority of neurons expressing calretinin in the submucosal plexus are S-100β positive, whereas only sparse calretinin- and nNOS-expressing myenteric neurons exhibit S100-β immunoreactivity, highlighting another major difference in the expression of S-100β between the two ENS plexi.

**Figure 7.**
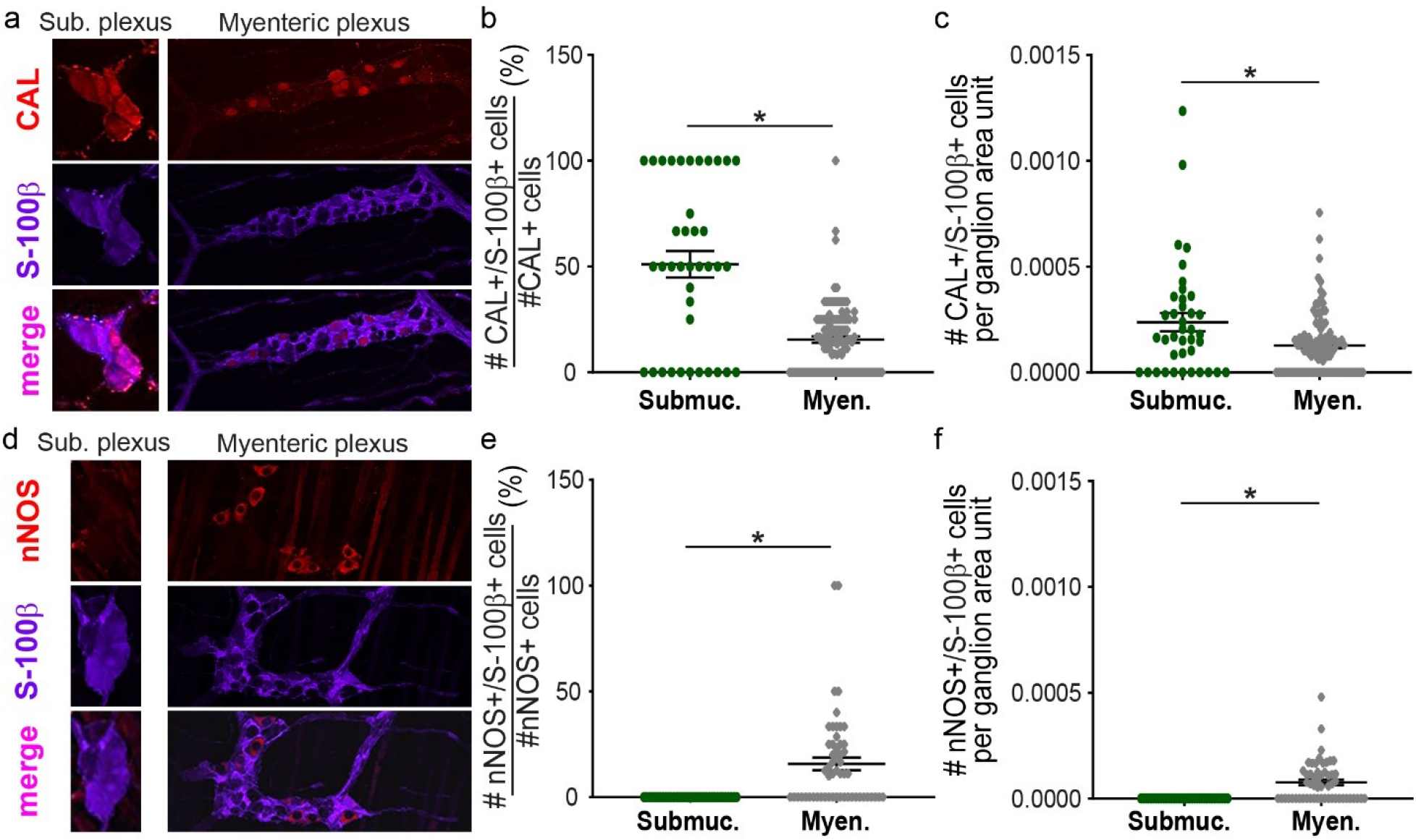
A majority of calretinin-expressing submucosal ganglionic cells are immunoreactive for S-100β, while only a minority of calretinin- and nNOS-expressing myenteric ganglionic cells are S-100β-positive. (a/d) Representative confocal images of ganglia from wholemount preparations of ileal submucosal and myenteric plexi from an adult mouse stained for calretinin (a) and nNOS (d) (red) and the EGC marker S-100β (magenta). (b/e) Quantification of the proportion of calretinin (CAL)-(b) or nNOS- (e) expressing cells immunoreactive for S-100β. Data are expressed as the percentages of the number of CAL^+^/ S-100β^+^ cells normalized to the total number of CAL^+^ cells per ganglion (b) and the percentages of the number of nNOS^+^/ S-100β^+^ cells normalized to the total number of nNOS^+^ cells per ganglion (e) (±SEM). (c/f) Quantification of the density of calretinin (CAL)-(b) or nNOS- (e) expressing cells immunoreactive for S-100β. Data are expressed as the number of CAL^+^/ S-100β^+^ cells normalized to ganglion area (c) and the number of nNOS^+^/ S-100β^+^ cells normalized to ganglion area (f) (±SEM). (b-c, e-f) Graphs show individual ganglia (n=35-109 ganglia) from 3 independent experiments. Mann-Whitney, *p<0.05.

## Discussion

Overlooked until the beginning of the twenty first century, EGC have recently drawn much attention, leading to compelling demonstration that EGC are potent regulators and/or effectors of a large majority of (if not all) GI functions, and that EGC defects are involved in several GI and extra-digestive diseases. Therefore, improving the current understanding of EGC biology may identify novel pathophysiological mechanisms and therapeutic targets. One specific gap of knowledge is the fairly poor appreciation of EGC heterogeneity. Here we report a simple and robust method to grow separately inEGC and outEGC from the same animal, allowing for the first time to directly interrogate phenotypical and functional differences between EGC from different layers of the small intestinal wall. Our data indicate that inEGC and outEGC primary cultures comprised distinct EGC subtypes and exhibit significantly different cellular response to purinergic signaling. We further highlight *ex vivo* differences in molecular identity between ganglionic EGC from inner plexus (submucosal) and outer plexus (myenteric). Altogether these studies provide direct evidence that EGC from different layers of the gut wall should be considered as distinct pools of EGC subtypes, and that our novel culture system is a highly valuable tool for future investigations thoroughly dissecting this cellular heterogeneity at the single cell level.

One important finding of our study is that inEGC and outEGC have distinct compositions on EGC subtypes based on (co-)expression of the three common EGC markers S-100β, GFAP and Sox10. Previous work by the Pachnis lab had reported differential (co-)expression of S-100β, GFAP and Sox10 in outEGC (myenteric EGC) subtypes^1^. Our work confirms these findings and demonstrates that such differential expression also exists in inEGC. Most importantly, our data show that the proportion of the distinct EGC subtypes -defined by different (co-)expression of S-100β, GFAP and Sox10-is different in inEGC and outEGC primary cultures. The most profound difference driving this observation was the differential expression of Sox10 between inEGC and outEGC primary cultures, with fewer inEGC expressing (lower levels) of Sox10 as compared to outEGC. We confirmed these findings *ex vivo* in ganglia of ENS wholemounts. These data are in agreement with a recent paper investigating age-differences in ENS neurochemichal coding showing that numbers of ganglionic Sox10-positive cells are much lower in the submucosal plexus when compared to the myenteric plexus in the mouse duodenum at 2 weeks and 6 weeks of age, although no statistics were reported as this was not the purpose of their investigation^19^. While we chose to use the level of expression of the three well-accepted EGC markers to define different EGC subtypes, it is very likely that these distinct molecular identities expand beyond expression level of EGC markers and dictate major differences in key regulatory pathways of EGC biology. This hypothesis seems particularly plausible considering that Sox10 is a transcription factor controlling glial lineage specification, differentiation and maintenance^21,22^. However interestingly our data indicate that inEGC and outEGC exhibit significantly different calcium response intensities to ATP stimulation, independently of their level of (co-)expression of S-100β, GFAP as well as Sox10. Indeed analyses of calcium responses to ATP at the single cell revealed that all inEGC, regardless of their level of (co-)expression of S-100β, GFAP and Sox10, showed more intense intracellular calcium changes when compared to corresponding outEGC subtype. These data suggest fundamental functional differences between EGC from different layers of the gut wall, and it would be extremely interesting in the future to identify pathways responsible for this differential sensitivity to ATP stimulation. Interestingly our statistical analyses did not reveal significant differences in calcium responses to ATP among the distinct inEGC subtypes or outEGC subtypes. This could appear in contradiction with the study by Boesmans and colleagues who reported that EGC present within myenteric ganglia (which was termed “Type I EGC”) showed significantly higher changes in intracellular calcium in response to ATP when compared to myenteric inter-ganglionic or extra-ganglionic EGC (“Type II EGC” and “Type III EGC”, respectively). However this putative discrepancy between the two studies may originate from the fact that different criteria were used to define the distinct EGC subtypes.

Another major finding of this study is the presence of large S-100β+/GFAP-ganglionic cells that express the two pan-neuronal markers PGP9.5 and HuC/D in both plexi, although, while this cell population represents more than half of PGP9.5+ and HuC/D+ cells in submucosal ganglia, it is significantly rarer in myenteric ganglia. Our data are in agreement with the paper by Parathan and colleagues who reported similar results at 2 weeks and 6 weeks of age in the mouse duodenum and colon, although they only focused on S-100β and HuC/D^19^. We also found that over half of calretinin-expressing submucosal ganglionic cells corresponded to this S-100β+/GFAP-/HuC/D+ or PGP9.5+ population. As calretinin-positive cells represent >90% of ganglionic cells in the submucosal plexus of mouse ileum^13^, this suggests that a large proportion, if not a majority of ganglionic cells are CAL+/S-100β+/GFAP- (and PGP9.5+/HuC/D+). As CAL+ neurons have reported to correspond to vasomotor and cholinergic and non-cholinergic secretomotor neurons in the submucosal plexus^13^, it would be very interesting to test whether CAL+ ganglionic cells expressing S-100β correspond to one specific type of neurons. In the myenteric plexus, the rare large S-100β+/GFAP-/HuC/D+ cells were found to correspond to nNOS+ or CAL+ cells, which together represent intrinsic primary afferent neurons (IPAN)(CAL+ and to a lesser extent nNOS+)^23,24^, descending (nNOS+) and ascending (CAL+) interneurons as well as inhibitory (nNOS+) and excitatory (CAL+) motor neurons^25^. Here also it would be very interesting to test whether CAL+ or nNOS+ ganglionic cells expressing S-100β correspond to one specific type of neurons.

Based on their size, ganglionic localization and morphology, the absence of expression of other EGC markers (GFAP) but the expression of pan-neuronal markers (PGP9.5 and HuC/D) and specific subtypes of neurons (CAL and nNOS), it is likely that these S-100β+/GFAP-/HuC/D+ or PGP9.5+ cells correspond to neurons, and not EGC. Thus, one can wonder S-100β role in these neurons and why it is mostly restricted to submucosal neurons. It is tempting to speculate that this may be in response to specific microenvironmental cues or -as suggested in previous paragraph-that S-100β expression is inherent to the specification and/or the function of a specific type of neurons. Also it is important to note that, as S-100β is frequently found in submucosal neurons, gross changes in S-100β level of expression in lysates of intestinal cross sections might not specifically reflect changes in EGC density.

Thus this work provides strong evidence that inEGC and outEGC are two distinct pools of EGC, and that a better understanding of their respective biological specificities may lead the identification of therapeutic targets to alleviate defects of discrete EGC involved in the regulation of (specific) mucosal functions or gut motility. To this effect, the novel culture method we report here to robustly grow heterogenous inEGC and outEGC cultures combined with wholemount preparations represent a powerful tool to investigate the respective roles of inEGC *versus* outEGC in various GI functions.

