## Supplemental Material for "Enteric glial cells of the two plexi of the enteric nervous system exhibit phenotypic and functional inter- and intra-heterogeneity"

### Primary antibodies

| Target | Host | Dilution | Supplier | Catalog number |
|---|---|---|---|---|
| Sox10 | goat | 1:500 | Santa Cruz | sc-17342 |
| S-100β | rabbit | 1:1,000 | Dako | Z031129-2 |
| GFAP | chicken | 1:500 | Abcam | ab4674 |
| PGP9.5 | chicken | 1:500 | Abcam | ab72910 |
| αSMA | mouse | 1:500 | Abcam | ab7817 |
| HuC/D biotin | mouse | 1:150 | Invitrogen | A-21272 |
| Calretinin | goat | 1:1,000 | Swant | CG1 |
| nNOS | sheep | 1:500 | Millipore | AB1529 |
| p75NTR | rabbit | 1:500 | Abcam | ab8874 |
| F4/80 | mouse | 1:300 | Invitrogen | MF48000 |

### Secondary antibodies

| Target | Fluorochrome | Host | Dilution | Supplier | Catalog number |
|---|---|---|---|---|---|
| goat IgG | AF568 | donkey | 1:500 | Invitrogen | A11057 |
| rabbit IgG | AF647 | goat | 1:500 | Invitrogen | A21245 |
| chicken IgG | AF488 | goat | 1:500 | Invitrogen | A11039 |
| rabbit IgG | AF568 | goat | 1:500 | Invitrogen | A10042 |
| mouse IgG | AF647 | donkey | 1:500 | Invitrogen | A31571 |
| rabbit IgG | AF488 | goat | 1:500 | Invitrogen | A11008 |
| streptavidin | AF568 | n/a | 1:500 | Invitrogen | S11226 |

**Supplemental Table 1. List of primary and secondary antibodies used for immunofluorescence studies.**

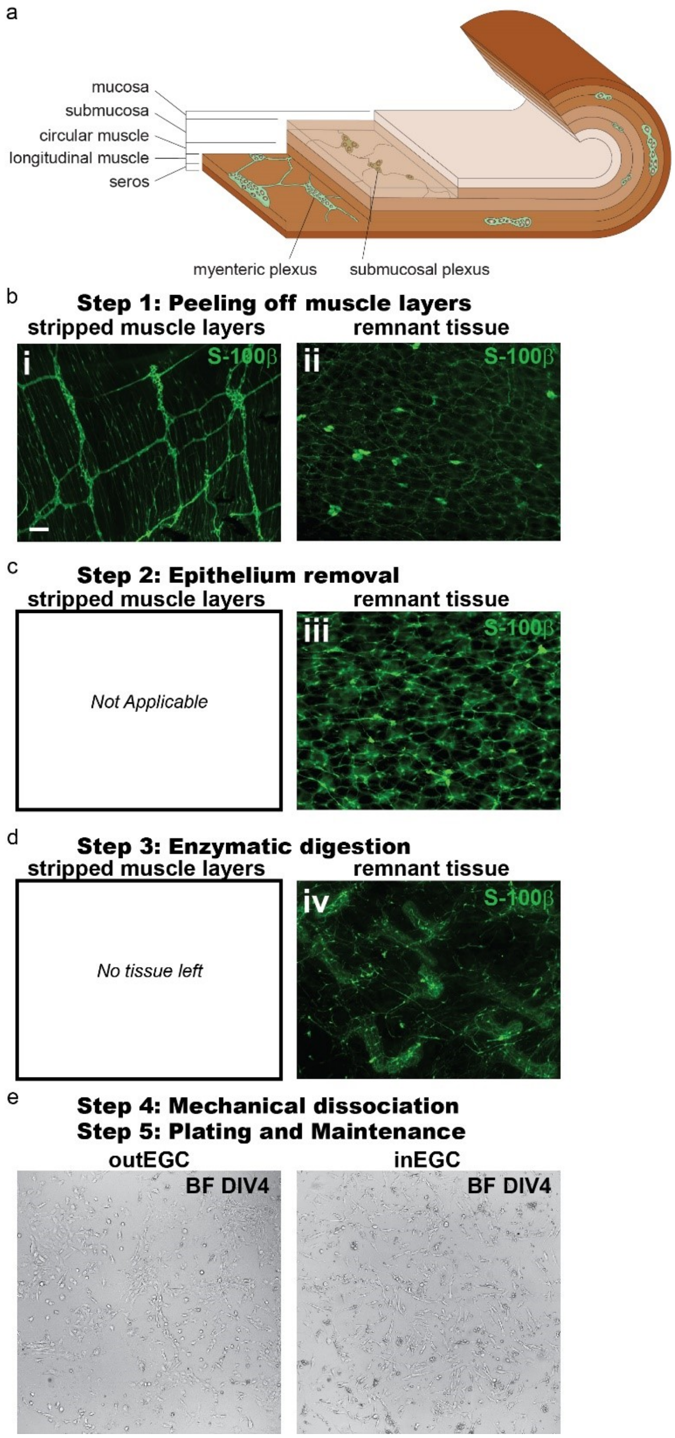


**Supplemental Figure 1. Outline of the methods used to isolate and establish primary cultures of inEGC and outEGC from the mouse small intestine.** (a) Schematic presenting the different layers of the small intestine wall. (b) Representative pictures of the muscle layers that were peeled off (i) and the remnant tissue (ii) stained for the EGC marker S-100β (green). (c) Representative pictures of the remnant tissue following EDTA/DTT treatment and mechanical dissociation to detach the epithelium (iii) stained for S-

100β (green). (d) Representative pictures following enzymatic digestion. Of note, no complete tissue was left following the digestion of the muscle strips. (e) Representative brightfield pictures of outEGC and inEGC primary cultures at DIV4.

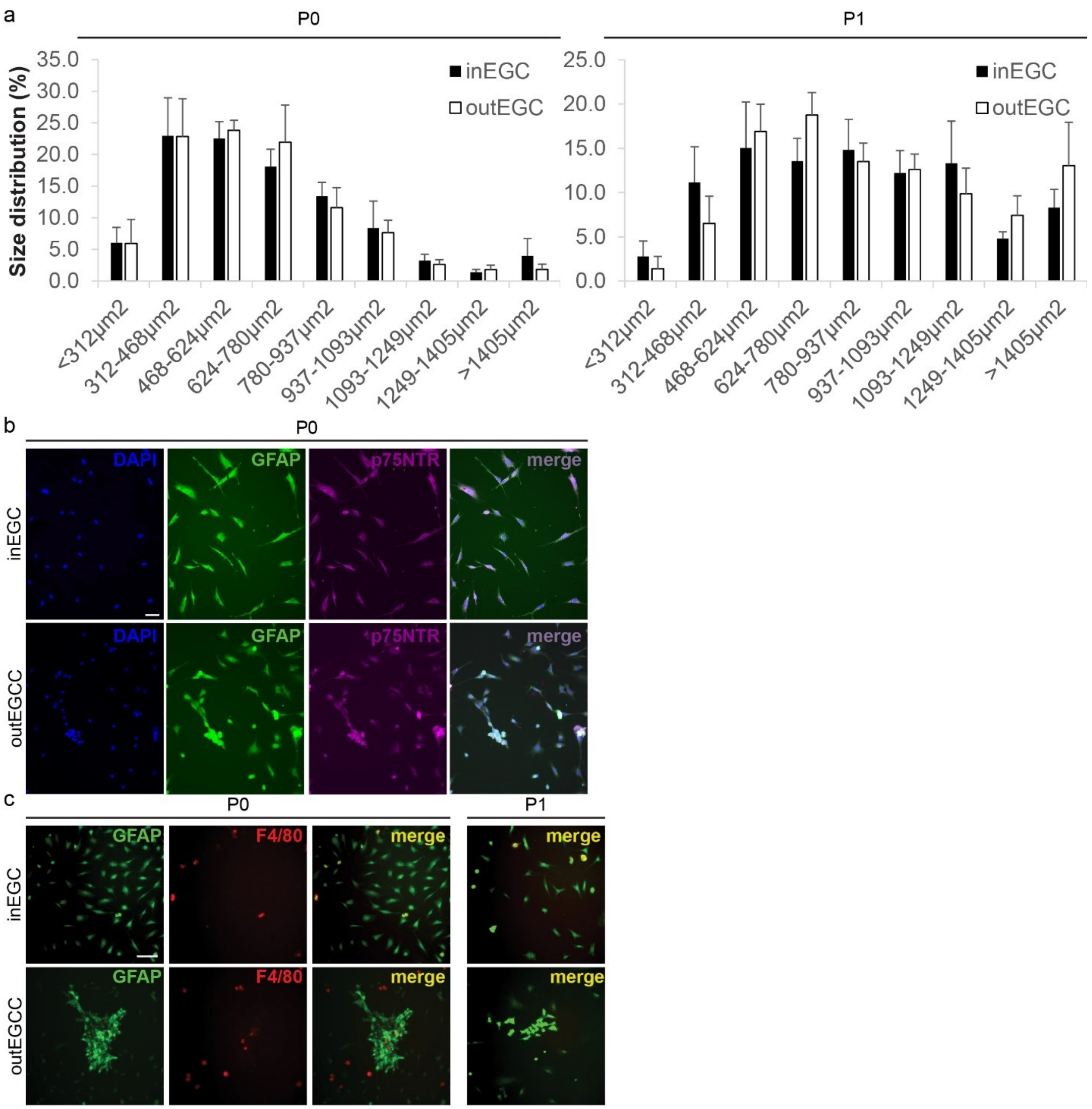


**Supplemental Figure 2. Primary cultures of inEGC and outEGC exhibit similar size distribution, are mostly composed of p75-expressing cells and very few F4/80 positive cells.** (a) Bar graphs representing the proportion of cells within different size ranges in P0 and P1 inEGC and outEGC primary cultures. Data are represented as the average of the percentages calculated per field (±SEM). 5.3±0.4 fields (61±3 cells) have been sized per condition. n=4 independent experiments. (b-c) Representative immunofluorescence pictures of P0 and P1 inEGC and outEGC cultures stained for p75 (b) (Magenta) and F4/80 (c) (red). EGC are marked with GFAP (green). Scale bars = 50 µm (b) and 100µm (c).

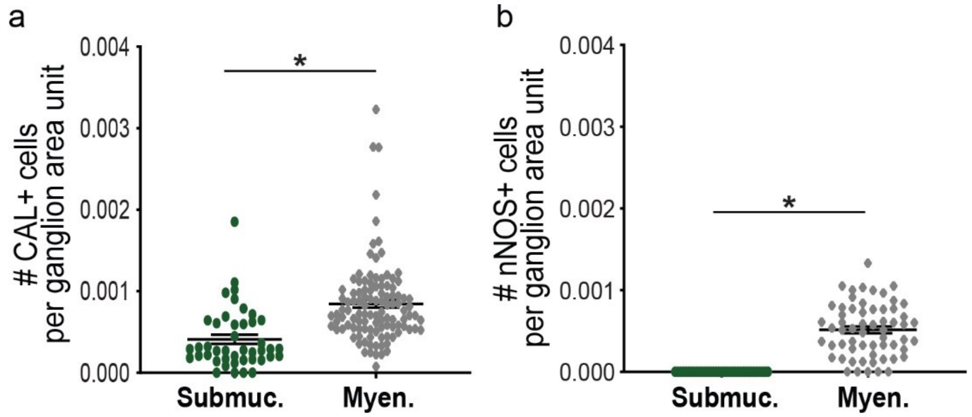


**Supplemental Figure 3. Density of calretinin and nNOS-expressing cells in submucosal and myenteric ganglia.** (a/b) Quantification of the density of calretinin (CAL)-(a) or nNOS- (b) expressing cells per ganglion area unit. Data are expressed as the number of $CAL^{+}$ cells normalized to ganglion area (a) and the number of $nNOS^{+}$ cells normalized to ganglion area (b) (±SEM). Graphs show individual ganglia (n=35-109 ganglia) from 3 independent experiments. Mann-Whitney, *$p<0.05$.
